# SPARC promotes insulin secretion through down-regulation of RGS4 protein in pancreatic β-cells

**DOI:** 10.1101/2020.01.11.902916

**Authors:** Li Hu, Fengli He, Meifeng Huang, Qian Zhao, Lamei Cheng, Neveen Said, Zhiguang Zhou, Feng Liu, Yan-Shan Dai

## Abstract

SPARC-deficient mice have been shown to exhibit impaired glucose tolerance and insulin secretion, but the underlying mechanism remains unknown. Here, we show that SPARC enhanced the promoting effect of Muscarinic receptor agonist oxotremorine-M on insulin secretion in cultured mouse islets. Overexpression of SPARC down-regulated RGS4, a negative regulator of β-cell M3 muscarinic receptors. Conversely, knockdown of SPARC up-regulated RGS4 in Min6 cells. RGS4 was up-regulated in islets from *sparc* -/- mice, which correlated with decreased glucose-stimulated insulin secretion (GSIS). Furthermore, inhibition of RGS4 restored GSIS in *sparc* -/- mice, and knockdown of RGS4 partially decreased the promoting effect of SPARC on oxotremorine-M stimulated insulin secretion. Phosphoinositide 3-kinase (PI3K) inhibitor LY-294002 abolished SPARC-induced down-regulation of RGS4. Taken together, our data revealed that SPARC promoted GSIS by inhibiting RGS4 in pancreatic β cells.

## Introduction

The etiology of Diabetes involves both insulin resistance and decreased insulin secretion in β-cells^1-3^. Adipose tissue inflammation plays an important role in the development of obesity-induced insulin resistance^2,4^. In the prediabetic insulin-resistant state, islets respond to the increased insulin demand by increasing insulin secretion and β-cell mass, which generates compensatory hyperinsulinemia and maintains relative normal glycaemia. When insulin secretion is insufficient to compensate for the insulin resistance, type 2 diabetes emerges. The early β cell dysfunction is manifested as impaired glucose-stimulated insulin secretion (GSIS), and loss of first phase GSIS can be detected in the earliest stages of type 2 diabetes^1,3^.

SPARC (Secreted Protein Acidic and Rich in Cysteine) is expressed mainly in bone and adipocytes, and it is a secreted extracellular matrix protein that is produced as a 32-kDa matricellular glycoprotein ^5-9^. SPARC mediates interactions between cells and their extracellular matrix, and targeted deletion of sparc results in compromised extracellular matrix in mice ^7,10^. SPARC can bind to integrin beta3 and activate integrin linked kinase ^11^. Secretion of SPARC by adipose tissue is increased by insulin and leptin ^12,13^. Adipose tissue is a major source of SPARC, which in turn inhibits adipogenesis and induces insulin resistance in adipose tissue in an autocrine or paracrine fashion^14-16^. SPARC has been shown to be involved in oxidative stress, neurogenesis, insulin resistance, glucose metabolism and Glut4 expression ^16-20^.

SPARC is expressed in the stromal cells of mouse primary islets that can be detected by Western blot, and plays a role in reducing IGF-1-induced islet survival ^21^. *sparc*-/- mice on high-fat diet showed an impaired insulin secretion capacity and a dramatically reduced expression of glucose transporter 2 in the islets ^22^. SPARC-deficient mice exhibited impaired glucose tolerance ^22,23^, and one of the underlying mechanism is reduced insulin secretion ^22^.

Acetylcholine, which is released from intrapancreatic nerve endings, stimulates insulin secretion through binding and activation of M3 muscarinic receptor, resulting in increased intracellular calcium concentration in beta-cells during food ingestion ^24^. Mice selectively lacking M3 muscarinic receptor in pancreatic β-cells displayed impaired glucose tolerance and a dramatic reduction in insulin secretion, demonstrating that M3 muscarinic receptor in β-cells is essential for maintaining normal glucose homeostasis ^24 25^. RGS4 is a negative regulator of G protein-coupled M3-muscarinic receptor. Pancreatic beta cell-specific knockout of Rgs4 did not alter basal glucose levels, but increased plasma insulin levels and muscarinic agonist-stimulated insulin release ^26^, suggesting that selectively enhancing the M3 muscarinic receptor signaling in pancreatic β-cells could lead to novel treatment for patients with type 2 diabetes ^26^.

To elucidate the mechanism by which SPARC regulates insulin secretion and β cell physiology, we had analyzed insulin secretion in SPARC-deficient β cells. Our results demonstrate that SPARC was essential for normal β cell function and promotes insulin secretion by inhibiting RGS4 expression.

## Materials and Methods

### Materials

Sparc -/- mice were a gift from Dr. Neveen Said at University of Virginia, Charlottesville, Virginia, USA ^27^ and were maintained in the accredited pathogen-free Second Xiangya hospital mice facility on a 12 h light/dark cycle ^28^. C57BL/6 mice were purchased from Model Animal Research Center of Nanjing University. All experiments and methods were performed following approval of the protocols by the Animal Care Research Committee of Second Xiangya Hospital. All methods were performed in accordance with the relevant guidelines and regulations.

Human sparc cDNA clone was described by us before ^29^ and subcloned into pShuttle vector (Clontech). Adenovirus expressing human SPARC was constructed using Adeno-X expression system (Clontech) as described before ^30,31^. GAPDH, Horseradish peroxidase labeled donkey anti rabbit or donkey anti mouse antibodies were from Cell Signaling (Beverly, MA). Recombinant mouse SPARC protein (cat. number 942-SP-050) was purchased from R & D systems. Oxotremorine M (Oxo-M) and LY294002 were purchased from Sigma-Aldrich. CCG-4986 was purchased from ChemBridge (San Diego, CA).

## Methods

### Cell Culture

Min6 cells were originally purchased from ATCC and were cultured in DMEM containing 15 % FBS, 25 mM Glucose and 50 uM β-mercaptoethanol as described before ^31^. Min6 cells were seeded in a six-well plate and allowed to attach overnight. Min6 cells were incubated for 24 h before collection and analysis.

### Islet isolation

Islets were isolated from 8 to 12 weeks old C57BL/6 male mice as described before by our laboratory^31-34^. Briefly, mouse islets were isolated using perfusion and digestion of pancreas with collagenase V (from Roche), density gradient purification with histopaque-1077 (Sigma), and then hand-picked. Isolated islets were cultured overnight in RPMI 1640 containing 10%FBS, 11 mM glucose, and then switched for 1 hr to Krebs Ringer Bicarbonate buffer containing 2.6 mmol/l CaCl2/2H2O, 1.2 mmol/l MgSO4/7H2O, 1.2 mmol/l KH2PO4, 4.9 mmol/l KCl, 98.5 mmol/l NaCl, and 25.9 mmol/l NaHCO3 (all from Sigma-Aldrich) supplemented with 20 mmol/l Na-HEPES and 0.1% BSA. About 10 islets in each experimental condition were transferred to each well in 24-well plate containing 2.8 mM and 16.7 mM glucose concentration in Krebs Ringer Bicarbonate buffer for 1 hr. The supernants were collected for insulin measurements. The islets were lysed with 1% Triton to determine total protein content in the islets. Insulin levels were measured with an ELISA kit from ALPCO. About 200 isolated mouse islets from WT or sparc ko mice were also collected for Western blot analysis of RGS4 and SPARC.

### Western blots

Western blots were performed using equal amounts of whole cell extract protein as describe before ^33,35^. Briefly, cell lysates were run on SDS PAGE, proteins transferred to a nitrocellulose membrane. The membranes were incubated with the primary antibodies, rabbit anti-RGS4 (Santa cruz), mouse anti-SPARC (Haematologic Technologies, Inc), rabbit anti-AKT-S473, rabbit anti-AKT, mouse anti-beta-actin (Cell Signaling), respectively, followed by Horseradish peroxidase labeled donkey anti rabbit or donkey anti mouse antibodies. Protein signal was visualized by using Immun-Star chemiluminescent kit (Bio-Rad) and quantified by Bio-Rad Imager.

### Adenoviral infection

Adenovirus was amplified in 293 HEK cells. The adenovirus in 293 cells was collected and subjected to three cycles of freeze-thaw. Adenovirus titer was determined by using Adeno-X rapid titer kit from Clontech. Min6 cells or isolated islets were infected with Ad-SPARC or Ad-EGFP at 100 MOI for 16 hours. The next day, the medium was changed and the cells were cultured for total 48 hours. We used Ad-GFP virus as control and the expression of GFP inside the core of islets was confirmed under fluorescent microscope.

### siRNA transfection

Min6 cells were transfected with *sparc* siRNA or control scramble siRNA using Metafectene Pro, a cationic liposome based transfection reagent (Biontex laboratories, Germany), which possessed high transfection efficiency in Min6 cells shown by the manufacturer. siRNA against mouse SPARC and mouse RGS4 were synthesized by Shanghai GenePharma Co. The sense sequence of mouse SPARC was: AGAAGCUGCGUGUGAAGAAUU. The sense sequence of mouse RGS4 was: CCUGCCUGAGGAGUCAAATT.

### RT-qPCR

RNA was first extracted using Trizol reagent (Invitrogen), then followed by purification and DNase I digestion to remove genomic DNA contamination using RNAeasy column (Qiagen) according to the manufacturer’s protocol. After reverse transcription (Thermo Scientific), cDNA were quantified with ABI 7900HT Fast Real-Time PCR System (Applied Biosystems). Relative gene expression levels were calculated using the ΔΔCt method, with 18S or GAPDH used as the reference gene and normalized. RGS4 qPCR primer sequences were: Sense: AGAAATGGGCTGAATCGTTG; Anti-sense: GGCTTGTCTCCTCTCTGGTG.

### Microarray method

Min6 cells were infected with Ad-EGFP or Ad-SPARC for 48 hours. The cells were collected and total RNA was prepared by Trizol and RNA easy column (Qiagen). The RNA samples were sent to Shanghai Biotechnology incorporation for customer cDNA microarray analysis using Affymetrix mouse array 430 2.0.

### Statistics

Statistical significance was assessed by two-tailed Student t test. For multiple groups, statistical significance was performed with one way or two way ANOVA followed by Tukey’s multiple comparisons test by GraphPad Prism 7. A value of P <0.05 was considered significance. All data are expressed as the mean +/- SE.

## Results

### SPARC enhances insulin secretion in mouse pancreatic β-cells

To determine the effect of SPARC on insulin secretion, we overexpressed SPARC by adenovirus in isolated mouse islets. SPARC overexpression enhanced muscarinic receptor agonist Oxotremorine M-stimulated insulin secretion (Fig. 1A). To confirm this effect, we also treated cells with recombinant SPARC protein. As shown in Fig. 1B, SPARC enhanced Oxotremorine M-stimulated insulin secretion in mouse islets.

**Fig. 1.**
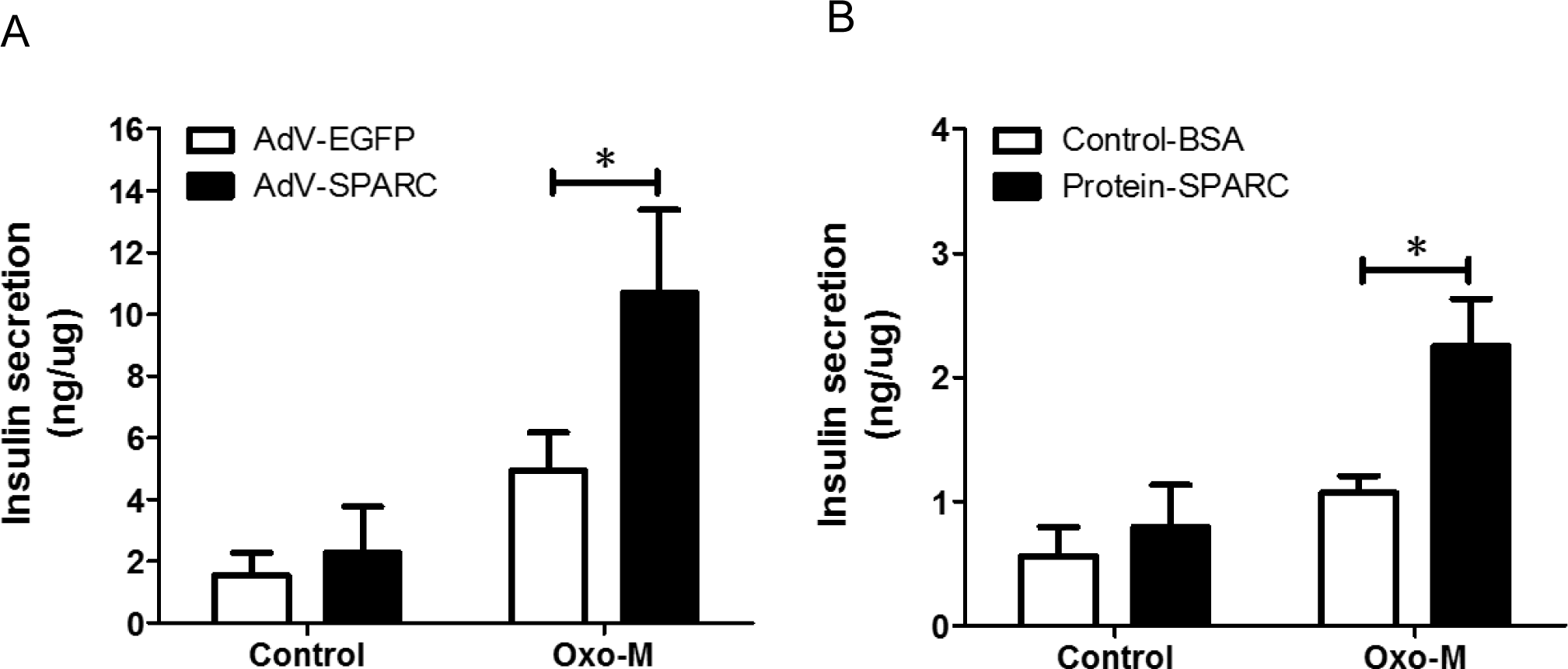
SPARC enhanced muscarinic receptor agonist-stimulated insulin secretion. (A) Mouse islets were isolated and culture in RPMI1640 containing 11 mM glucose overnight. The islets were infected with Ad-GFP or Ad-SPARC for 24 hrs. The islets were stimulated with 16.7 mM glucose, or 16.7 mM glucose and 100 µM Oxo-M in Krebs Ringer Bicarbonate buffer for 1 hr. The supernatants were collected and insulin was determined with insulin ELISA kit. The results were derived from three individual experiments. * denotes P<0.05, n=3. (B) Mouse islets were cultured in RMPI1640 medium with 11 mM glucose. Groups of islets were then incubated with Krebs Ringer Bicarbonate buffer and with 2.8 mM glucose, or 16.7 mM glucose in the presence or absence of recombinant SPARC, 1 µg/ml, or 16.7 mM glucose and 100 µM Oxo-M in the presence or absence of SPARC for 1 hr. * denotes P<0.05, n=3. Two-way ANOVA followed by Tukey’s multiple comparisons test was used in statistical analysis.

### Downregulation of RGS4 by Overexpression of SPARC

To study the changes of gene expression induced by overexpression of SPARC, we conducted cDNA microarray in Min6 cells overexpressing SPARC using Affimetrix mouse cDNA chip and found down-regulation by SPARC of RGS-4 mRNA to 0.38-fold compared to Ad-EGFP-infected Min6 cells (Fig. 2A); SPARC also induced MafA, Glut2, Notch2 and NeuroD2 by >2-folds as shown in Fig. 2A.

**Fig. 2.**
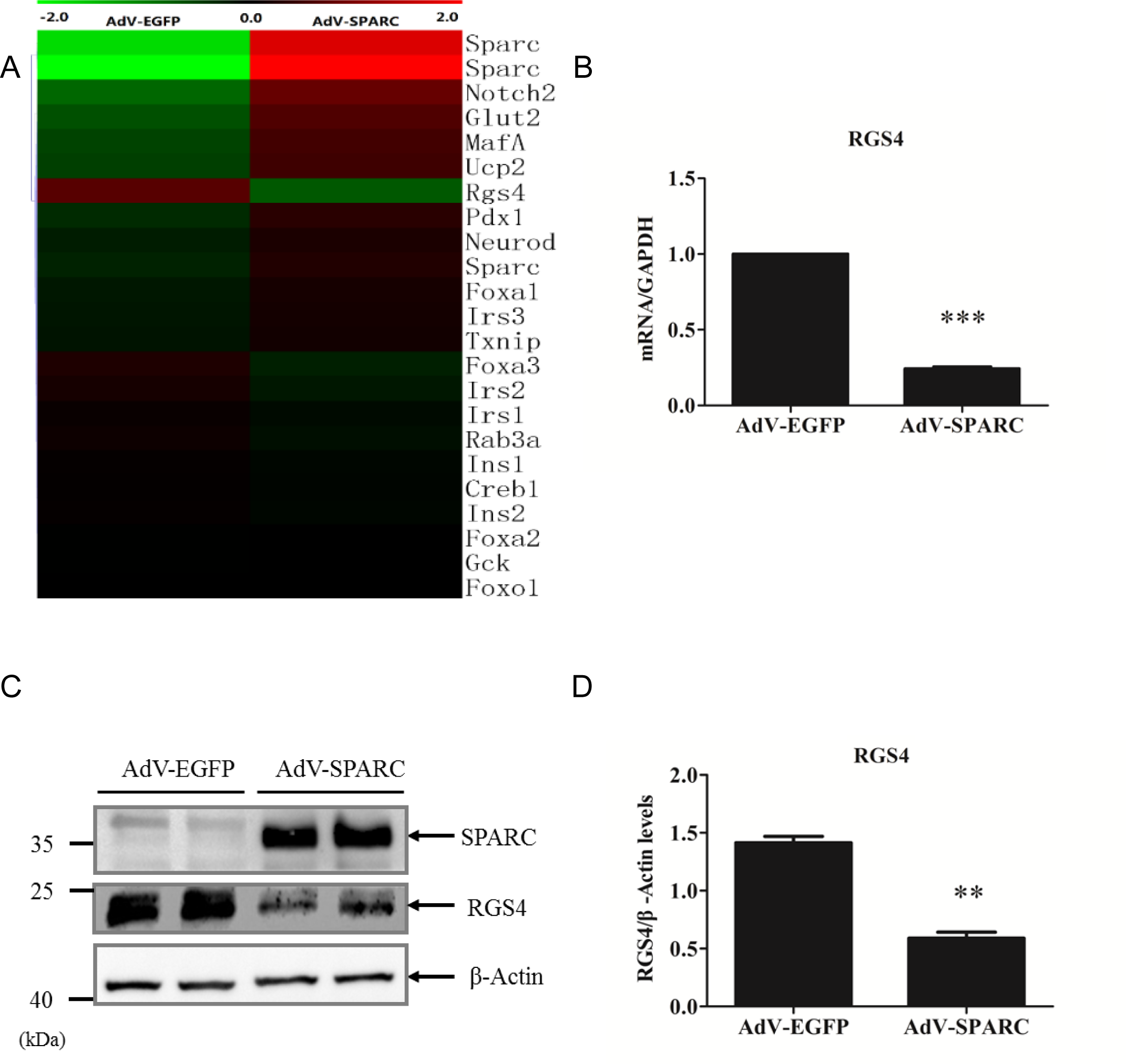
Overexpression of SPARC down-regulates RGS4 mRNA and protein in Min6 cells. Min6 cells were infected with Ad-EGFP or Ad-SPARC for 24 hrs. (A) mRNA were prepared from Ad-EGFP and Ad-SPACR treated Min6 cells and were subject to cDNA microarray; Microarray data measuring relative expression of selected genes were shown in the heat map. (B) RGS4 mRNA was down-regulated by overexpression of SPARC in Min6 cells. The cells were harvested for RNA preparation. RT-qPCR was performed to determine RGS4 mRNA levels after SPARC overexpression. (C) The cell lysates were subject to Western blotting with anti-SPARC antibody or anti-β-actin antibody and SPARC protein was shown to be overexpressed. The cell lysates were also subject to Western blotting with anti-RGS4 antibody and the results showed down-regulation of RGS4 protein by overexpression of SPARC in Min6 cells. (D) Relative RGS4 protein levels were quantified and normalized with β-actin. Statistical significance was assessed by unpaired two-tailed Student’s t test. ** denotes P<0.01, n=3. *** denotes P<0.001, n=3.

Since RGS4 is a known negative regulator of insulin secretion through inhibition of muscarinic M3 receptor-G(q) protein coupling in β cells, we further investigated the effect of SPARC overexpression and loss of *sparc* on RGS4 expression. To confirm the microarray data, we overexpressed SPARC and analyzed RGS4 mRNA levels by RT-qPCR. As shown in Fig. 2C, SPARC was overexpressed in Min6 cells by adenovirus-mediated transduction. Overexpression of SPARC in Min6 cells induced down-regulation of RGS4 mRNA (Fig. 2B) and protein (Fig. 2C, D).

### Up-regulated RGS4 protein in pancreatic islets in sparc -/- mice

We reasoned that if SPARC inhibits RGS4 expression, knockdown of SPARC would increase RGS4 expression. As shown in Fig. 3A, treatment with siRNA against *sparc* gene transcripts reduced SPARC expression by 60% in Min6 cells (Fig. 3 A) and the reduction of SPARC resulted in the increase in RGS4 expression by 2.5-fold (Fig. 3A, B).

**Fig. 3.**
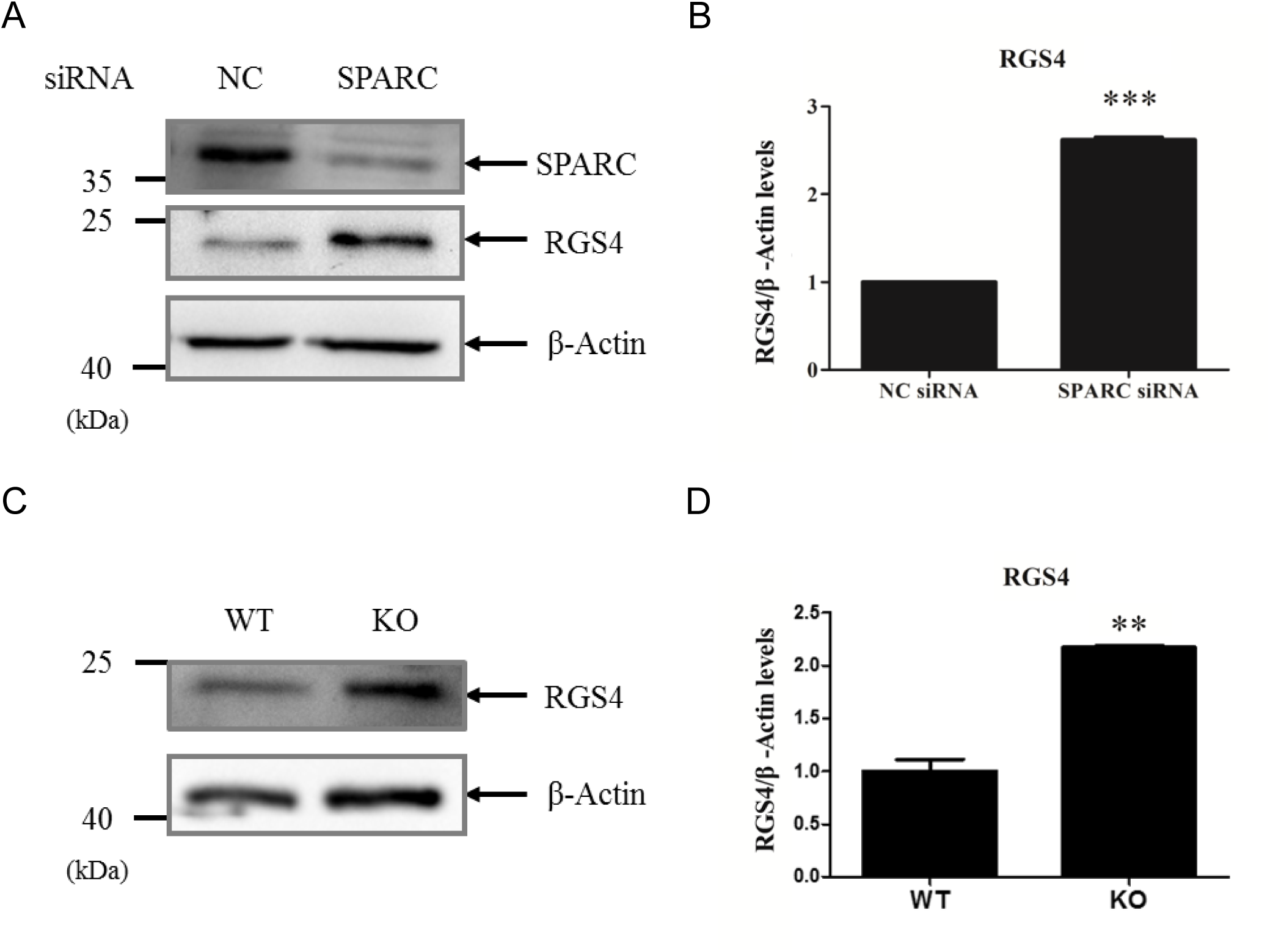
Knocking down of SPARC with siRNA in Min6 cells up-regulated RGS4 mRNA, and loss of *sparc* up-regulated RGS4 protein in isolated islets. Min6 cells were transfected with SPARC siRNA and cultured for 48 hrs. (A, B) The cell lysates were subject to Western blotting with anti-SPARC antibody or anti-beta-actin antibody (A). (B) Relative RGS4 levels were quantified and normalized with beta-actin. *** denotes P<0.001, n=3. Statistical significance was assessed by unpaired two-tailed Student’s t test. (C, D) Up-regulation of RGS4 protein in the islets of *sparc* -/- mice. Mouse islets were isolated from WT or *sparc* -/- mice and subject to Western blot analysis of RGS4 protein (C). Average RGS4 levels from two experiments were quantified and normalized to β-actin levels (D). ** denotes P<0.01, n=3.

To further corroborate the siRNA result in Fig. 3A and B, we also examined the effect of loss of *sparc* gene on expression of RGS4 protein in the islet of *sparc* -/- mice. We isolated islets from wild type (WT) and *sparc* -/- mice, performed Western blot analysis of RGS4 on these islets, and found that RGS4 protein levels were also up-regulated in the islets of *sparc* -/- mice (Fig. 3C, D).

### SPARC down-regulates RGS4 partially through activating Phosphoinositide 3-kinase and AKT signaling pathway

Since PI3K/Akt/GSK3beta pathway reduces IL-1beta-induced upregulation of RGS4 expression by inhibiting NF-kappaB activation^36^, we hypothesized that SPARC down-regulates RGS4 through PI3K/Akt pathway. The results showed that overexpression of SPARC in Min6 cells induced AKT 473 phosphorylation (Fig. 4A, C), whereas knockdown of SPARC reduced AKT 473 phosphorylation (Fig. 4B, D).

**Fig. 4.**
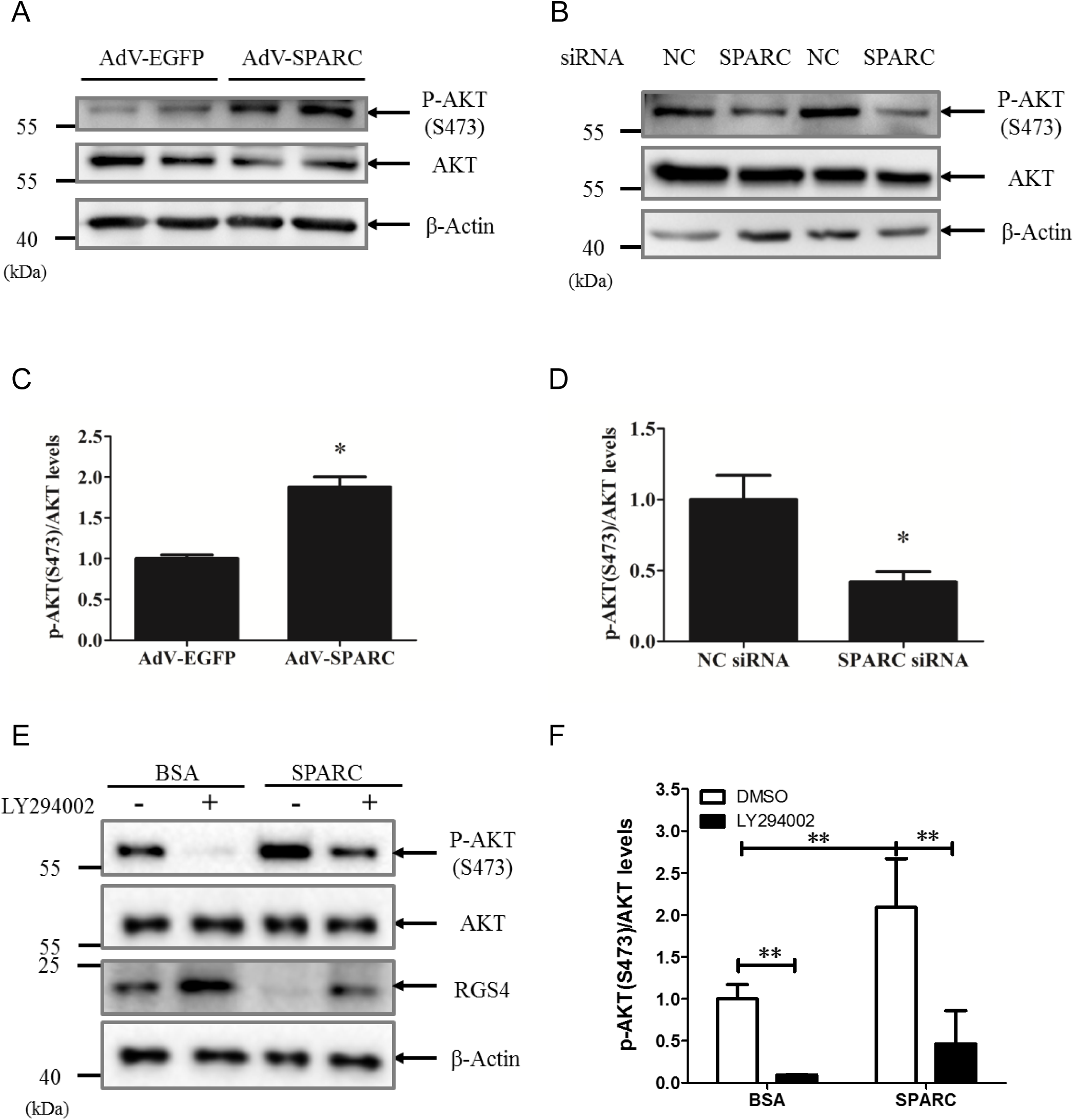
SPARC-induced RGS4 down-regulation in Min6 cells is abolished by PI3 kinase inhibitor LY-294002. (A, C) Overexpression of Ad-SPARC in Min6 cells increased AKT S473 phosphorylation. Min6 cells were infected with Ad-EGFP or Ad-SPARC for 24 hrs. The cell lysates were subject to Western blotting with anti-p-AKT (473) or anti-β-actin antibody. Lower panel was ratio of p-AKT S473 /total AKT quantified from three experiments (C). (B, D) Knocking down of SPARC in Min6 cells decreased AKT S473 phosphorylation. Min6 cells were transfected with SPARC siRNA and cultured for 48 hrs. The cell lysates were subject to Western blotting with anti-p-AKT(S473) antibody or anti-beta-actin antibody (B). Lower panel was ratio of p-AKT S473 /total AKT quantified from three experiments (D). (E, F) SPARC-induced RGS4 downregulation was blocked by PI3K inhibitor. Cultured Min6 cells after serum starvation for 12 hrs were pretreated with the PI3K inhibitor LY-294002 (10 μM) for 1 hr before treatment with or without recombinant SPARC (1 µg/ml) for 12 hrs followed by Western blot analysis with the indicated specific antibodies. Down-regulation of RGS4 by SPARC protein was abolished by LY-294002 (E). Lower panel was ratio of p-AKT S473 /total AKT quantified from three experiments (F). * denotes P<0.05, n=3. ** denotes P<0.01, n=3. Statistical significance was assessed by unpaired two-tailed Student’s t test.

To understand if SPARC down-regulates RGS4 through PI3K, we used LY294002, a selective inhibitor of PI3K, to treat Min6 cells. Consistent with the results in Fig.2C and Fig. 4A, the treatment of Min6 cells with recombinant SPARC protein induced AKT 473 phosphorylation and down-regulation of RGS4 protein. LY-294002 at 10 μM significantly decreased p-AKT (S473) expression levels and up-regulated RGS4 protein levels in Min6 cells (Fig. 4E, F). Furthermore, LY-294002 abolished the induction of AKT 473 phosphorylation and down-regulation of RGS4 protein by SPARC (Fig. 4E, F). Meanwhile, the changes of total AKT protein levels were not significant in either treatment. The data suggest that the activation of PI3K by SPARC played a role in the down-regulation of RGS4 expression. Taken together, these results indicated that in Min6 cells, SPARC down-regulated RGS4 expression levels at least partially through activating PI3K signaling pathway.

### Impaired insulin secretion in isolated islets of *sparc* -/- mice

To directly assess effect of SPARC on β-cell function, we carried out in vitro culture systems on isolated islets. First, to directly test the function of SPARC in β-cells, we measured insulin secretion under static incubation conditions by isolated islets obtained from WT and *sparc* -/- mice. As seen in Fig. 5A, the islets isolated from *sparc* -/- mice demonstrated significantly lower insulin secretion compared to those of WT mice in response to both high glucose and Oxo-M stimulation. Taken together, these data indicated that SPARC was necessary for normal insulin secretory function in response to glucose and Oxo-M in vitro in mouse β-cells.

**Fig. 5.**
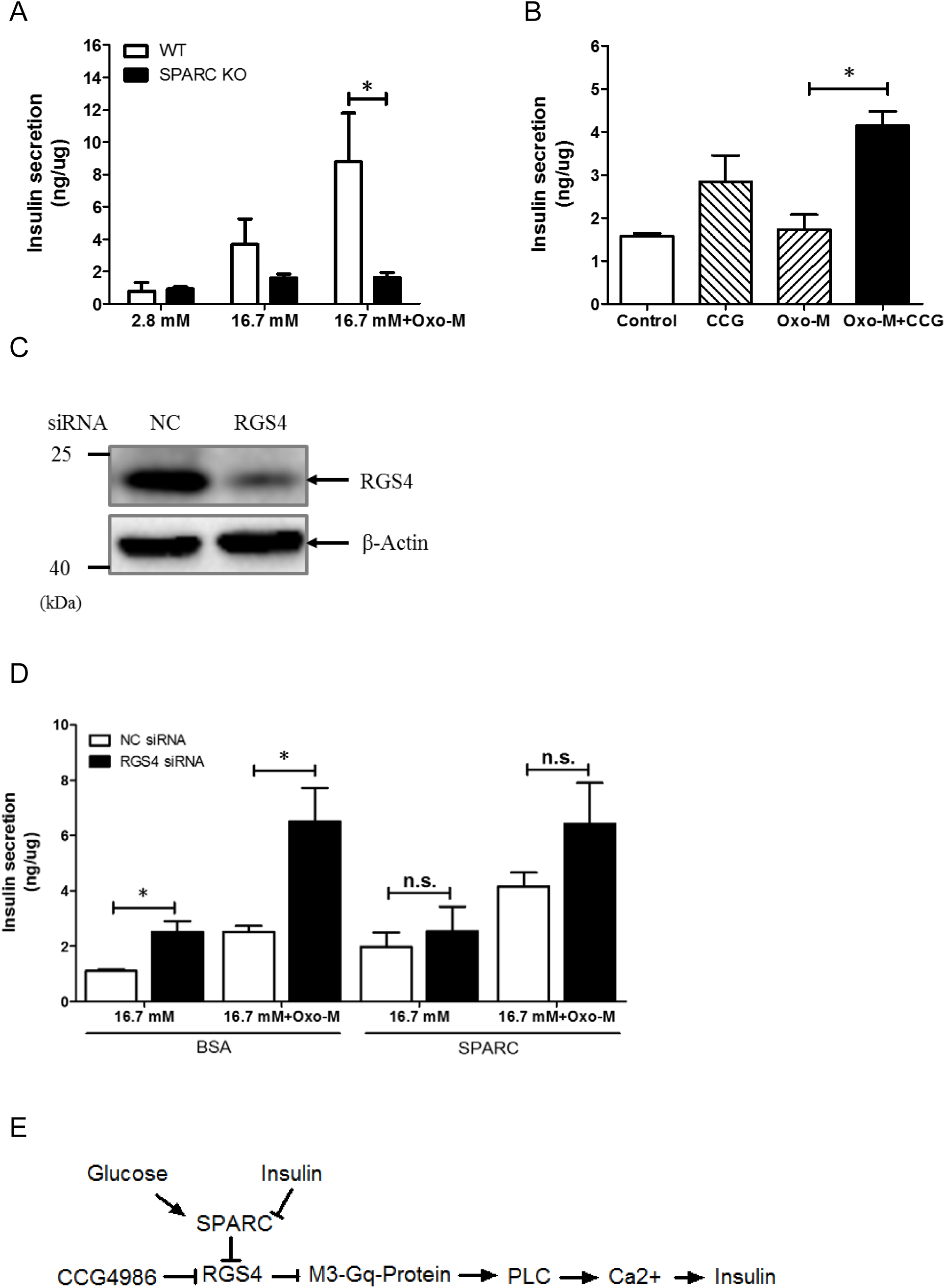
RGS4 inhibitor CGG-4986 restored insulin secretion in islets of *sparc* -/-mice. (A) Mouse islets were isolated from WT or *sparc* -/- mice. The isolated islets were cultured for 16 hrs in RPMI1640 with 11 mM Glucose and 10% FBS. The WT or *sparc* -/- islets were incubated for 1 hr in 2.8 mM glucose, then the medium was changed to 16.7 mM glucose, 16.7 mM glucose with Oxo-M, 100 µM in Krebs Ringer Bicarbonate buffer for 1 hour. The supernatants were collected for insulin assay. Shown were the levels of insulin secretion normalized to total protein concentrations. * denotes P<0.05, n=3. Two-way ANOVA followed by Tukey’s multiple comparisons test was used in statistical analysis. (B) *sparc* -/- mouse islets were incubated for 1 hr in 2.8 mM glucose, and then the medium was changed to 16.7 mM glucose, 16.7 mM glucose with Oxo-M, 100 µM, or 16.7 mM glucose + CGG 4986, 100 µM, or 16.7mM glucose, Oxo-M, 100 µM, CGG4986, 100 µM in Krebs Ringer Bicarbonate buffer. * denotes P<0.05, n=3. Shown were the levels of insulin secretion normalized to protein concentration. Two-way ANOVA followed by Tukey’s multiple comparisons test was used in statistical analysis. (C, D) Min6 cells were transfected with RGS4 siRNA and cultured for 48 hrs. The cell lysates were subject to Western blotting with anti-RGS4 antibody or anti-beta-actin antibody (C). (D) After transfected with RGS4 siRNA for 48 hrs, groups of Min6 cells were then incubated with Krebs Ringer Bicarbonate buffer with 16.7 mM glucose, or 16.7 mM glucose and 100 µM Oxo-M in the presence or absence of recombinant SPARC, 1 µg/ml for 1 hr. * denotes P<0.05, n=3. Two-way ANOVA followed by Tukey’s multiple comparisons test was used in statistical analysis. (E) The graph schematically outlined a potential novel mechanism for the regulation of insulin secretion by SPARC. Increased SPARC inhibited RGS4 expression, the latter of which in turn would result in increased M3 receptor / G-protein coupling, leading to increased insulin secretion in β-cells.

### Inhibition of RGS4 in cultured islets from *sparc* -/- mice restores Oxo-M stimulated GSIS

To demonstrate that increased RGS4 in *sparc* -/- mice is directly linked to impaired GSIS, we inhibited RGS4 function by administering CCG4986, a specific RGS4 inhibitor, in cultured islets isolated from *sparc* -/- mice. In islets of *sparc* -/- mice, both 16.7 mM high glucose and high glucose plus Oxo-M agonist did not induce GSIS (Fig. 5A). However, RGS4 inhibitor CCG-4986 restored high glucose and Oxo-M-induced GSIS (Fig. 5B). CCG-4986 increased insulin secretion by 3-fold, compared with control DMSO under high glucose plus Oxo-M condition. Furthermore, in order to confirm that SPARC regulates insulin secretion principally through RGS4, we used siRNA against *rgs4* gene transcripts to knockdown RGS4 expression. As shown in Fig. 5C, treatment with siRNA reduced RGS4 expression by 80% in Min6 cells (Fig. 5 C), and knockdown of RGS4 partially decreased the promoting effect of SPARC on oxotremorine-M stimulated insulin secretion (Fig. 5D). Taken together, these results suggested a potential pathway by which SPARC regulated insulin secretion through RGS4 (Fig. 5E).

## Discussion

In this study, we found that overexpression of SPARC inhibited RGS4 expression in Min6 cells; conversely, absence of SPARC increased RGS4 expression in the islets of sparc -/- mice. SPARC overexpression or exogenously administered recombinant SPARC increased muscarinic receptor agonist-stimulated insulin secretion. Here, we describe a novel mechanism for regulating β-cell function through SPARC-RGS4 pathway (Fig. 5E). The impaired insulin secretion represents a primary β-cell defect because isolated islets from sparc mice exhibited impaired GSIS compared to WT islets.

Previous study showed that SPARC increases human islet insulin secretion ^37^. Stable SPARC-expressing rat INS cell line also increases high glucose stimulated insulin secretion, but the mechanism was not elucidated ^37^. Our study in Min6 cell line and in isolated mouse islets showed that SPARC increased muscarinic receptor agonist-stimulated insulin secretion, suggesting that one of the mechanisms by which SPARC increases insulin secretion may be through suppression of RGS4 expression. RGS4 is one of seven members of the classic R4 RGS protein family that accelerates the intrinsic GTPase activity of Gαi/o and Gαq/11 family members. RGS4 inhibits the strength and duration of Gq signaling. In β-cells, RGS4 has been shown to negatively regulate insulin secretion through regulation of M3 receptor-G protein coupling^26^. During meals, acetylcholine is released from intra-pancreas parasympathetic nerve endings and bound to M3 receptors in β-cells, resulting in activation of PLCγ, IP3 and calcium release, and insulin secretion. We showed that increased SPARC expression inhibited RGS4 expression, which in turn would result in increased M3 receptor G-protein coupling and increased insulin secretion (Fig. 5E).

Our results are consistent with a recent study demonstrating that SPARC is necessary for insulin secretion in mouse β cells and that *sparc* -/- mice displayed glucose intolerance and diabetes in an age-dependent manner^22^. Moreover, Glucose transporter 2 (Glut2) expression in β cells is also reduced in *sparc* -/- mice ^22^, which is in agreement with our cDNA microarray data showing induction of Glut2 by SPARC in Min6 cells (Fig. 2A). In mouse β cells, receptor-activated Gq leads to the sequential activation of ERK1/2 and IRS2 signaling and selective increases in the expression of islet genes involved in insulin synthesis (Ins2, Pcks1, and Pcks2), and Glut2, thus improving β cell function ^38^. In Rat INS1-M3 insulinoma cell line, M3 receptor–mediated activation of Gq leads to increased Irs2 expression via sequential activation of PLC (PLCβ), PKC, and ERK1/2 ^38,39^.

PI3K activation inhibits RGS4 mRNA and protein in rabbit colonic smooth muscle cells, which was reversed by PI3K inhibitor LY-294002^36,40^. We hypothesized that SPARC down-regulates RGS-4 expression through PI3K activation. Treatment of Min6 cells with LY-294002 blocked SPARC-induced down-regulation of RGS4 protein (Fig. 4E), demonstrating that PI3K pathway regulated RGS4 downregulation induced by SPARC in Min6 cells. Our study showed that SPARC increased AKT phosphorylation in Min6 cells (Fig. 4A), which is downstream of PI3K. Akt phosphorylation has also contributed to increased insulin secretion ^41^. SPARC also activates the PI3K/AKT pathway to promote proliferation and metastasis in oral squamous cell carcinoma ^42^, suggesting this pathway to be ubiquitous.

Our study suggested that it may be possible to use RGS4 inhibitors to increase insulin secretion. There are several RGS4 inhibitors that would be of interest to test if they can be used to increase insulin secretion. CCG-4986 inhibits RGS4 function through the covalent modification of two spatially distinct cysteine residues on RGS4 ^43^. Our in vitro GSIS experiments showed that CCG-4986 restored GSIS in the isolated islets of sparc -/- mice, suggesting that the potential mechanism of defective insulin secretion was due to upregulated RGS4.

RGS4 dysfunction has been linked to many diseases, including hypertension, schizophrenia, and Parkinson’s disease ^44-46^. RGS4 is induced by IL-1β through activation of NF-κB, and RGS4 induction can in turn cause β-cell dysfunction by inhibiting insulin secretory machinery as exemplified before ^36^. Our results suggested that increased RGS4 expression in *sparc* -/- β-cells was mechanistically linked to a more chronic state of β-cell dysfunction. Since blood SPARC and insulin levels are increased in obesity and early type 2 diabetes ^47^, our results also suggested the involvement of SPARC in islet adaptation to insulin resistance in humans by increasing muscarinic receptor-mediated insulin secretion. However, whether SPARC also downregulates RGS4 expression in human β-cells remains to be confirmed.

In conclusion, we had identified a novel mechanism for SPARC to regulate insulin secretion by down-regulation of RGS4 expression and showed that SPARC played a role in the physiology of insulin secretion and in G-protein signaling in β-cells. In the future, it will be interesting to learn whether SPARC can improve insulin secretion in patients with type 2 diabetes through RGS4 pathway.

## Acknowledgements

This work is supported by Natural Science Foundation of China grant no.81270905 (Y-S Dai).

## Author Contributions

L.H., F.H., M.H. researched data. Q.Z., L.C., N.S. contributed materials. F.L.and Z.Z. contributed to discussion, reviewed manuscript. L.H and Y-S.D. researched data, wrote manuscript.

Y-S.D. is currently employed by Bristol-Myers Squibb Company. The rest of authors declare that there is no conflict of interest that could be perceived as prejudicing the impartiality of the research reported.

